# High-resolution binding data of TFIID and cofactors show promoter-specific differences *in vivo*

**DOI:** 10.64898/2025.12.22.691305

**Authors:** Sergio G-M Alcantara, Simon Bourdareau, Melanie Weilert, Julia Zeitlinger

## Abstract

TFIID is instrumental in recognizing promoter sequences and initiating transcription, yet a cohesive understanding of how this complex interacts with and functions at different promoter types *in vivo* is still lacking. Here, we employed ChIP-nexus to capture high-resolution binding footprints of all *Drosophila* TFIID subunits across the genome. These footprints reveal TFIID sub-modules whose DNA contacts suggest new structural details. At different promoter types, the footprints of the TAFs are very similar, suggesting the presence of engaged TFIID across all promoters. In contrast, the binding profile of TBP is promoter-specific, enabling us to identify TATA, DPR, and TCT/housekeeping promoters *de novo*, along with their underlying core promoter elements. Notably, our data point to NC2 being specific for TBP binding at the TATA box and suggest that TATA promoters show both TAF-dependent and TAF-independent initiation *in vivo*. These data suggest a model for the increased burst size observed at TATA promoters and provide a comprehensive resource for linking structural and biochemical results to *in vivo* data.

---

TFIID is a multi-subunit complex crucial for recognizing core promoter sequences, assembling the pre-initiation complex (PIC), and initiating RNA polymerase II (Pol II) transcription at transcription start sites (TSSs)^1^. TFIID is composed of TATA-binding protein (TBP) and 13 TBP-associated factors (TAFs), together forming a three-lobed structure when reconstituted *in vitro^2^*. As revealed by elegant cryo-EM studies, these lobes become reorganized when TFIID binds the promoter^3–6^. One lobe of TFIID binds to the downstream region (a.k.a. lobe C), while another lobe engages the upstream core promoter region (a.k.a. lobe B) together with the third lobe (a.k.a. lobe A), which is more flexible and structurally more difficult to characterize^5,6^. During this reorganization, TFIID places TBP near the site of transcriptional initiation, which is likely a critical step for PIC formation^5–9^.

Despite remarkable advances, TFIID is best understood *in vitro*, while its role *in vivo* across various promoter types remains incompletely understood. Different promoter types produce transcripts with distinct bursting properties^10–12^. Notably, TATA promoters show efficient reinitiation resulting in transcription bursts with more transcripts than TATA-less promoters. Cryo-EM structures suggest that various promoter types are bound by engaged holo-TFIID^6^, but whether TFIID plays different functional roles across promoter types is not understood.

Since TFIID is very flexible *in vitro* and sometimes found in sub-stoichiometric compositions^13–17^, it is possible that TFIID preferentially binds some promoter types, adopts different conformations, or forms distinct sub-complexes. Alternatively, TFIID could influence the transcription initiation pattern by differentially mobilizing TBP. TBP’s contacts appear to be highly regulated since it is bound by several TAFs (TAF1, TAF11/13)^8^, as well as by NC2 and Mot1^18–24^. These observations raise the possibility that TFIID binding differs across promoter types, an idea that remains untested *in vivo*.

To investigate TFIID function *in vivo*, we performed high-resolution ChIP-nexus experiments to map the occupancy of all 14 TFIID subunits across the *Drosophila* genome, in which core promoters are well characterized^25^. Our data reveal that while TFIID’s modular structure is broadly conserved across promoters, TBP and NC2 display striking promoter-specific binding patterns. Notably, TBP’s pattern at TATA promoters suggests an additional TAF-independent mode of initiation, which we propose contributes to the large transcriptional burst size of TATA promoters.

## Results

### ChIP-nexus captures all TFIID subunits at promoters

To robustly detect the patterns by which each TFIID subunit binds to core promoters *in vivo* at the highest resolution, we used ChIP-nexus^26^. ChIP-nexus is a chromatin immunoprecipitation coupled to exonuclease (ChIP-exo) technique, where a 5’ exonuclease digests the DNA of formaldehyde-fixed chromatin until encountering a protein-DNA crosslink. The DNA libraries produced from pulling down each TFIID subunit show precise strand-specific footprints at the sites where the TFIID subunit is directly or indirectly crosslinked to DNA (Figure 1A). We performed ChIP-nexus for each TFIID subunit in *Drosophila* Kc167 cells under normal conditions (DMSO control), and after treatment with triptolide (TRI), which blocks transcription at the PIC stage and yields higher occupancy of TFIID^27–30^.

**Figure 1.**
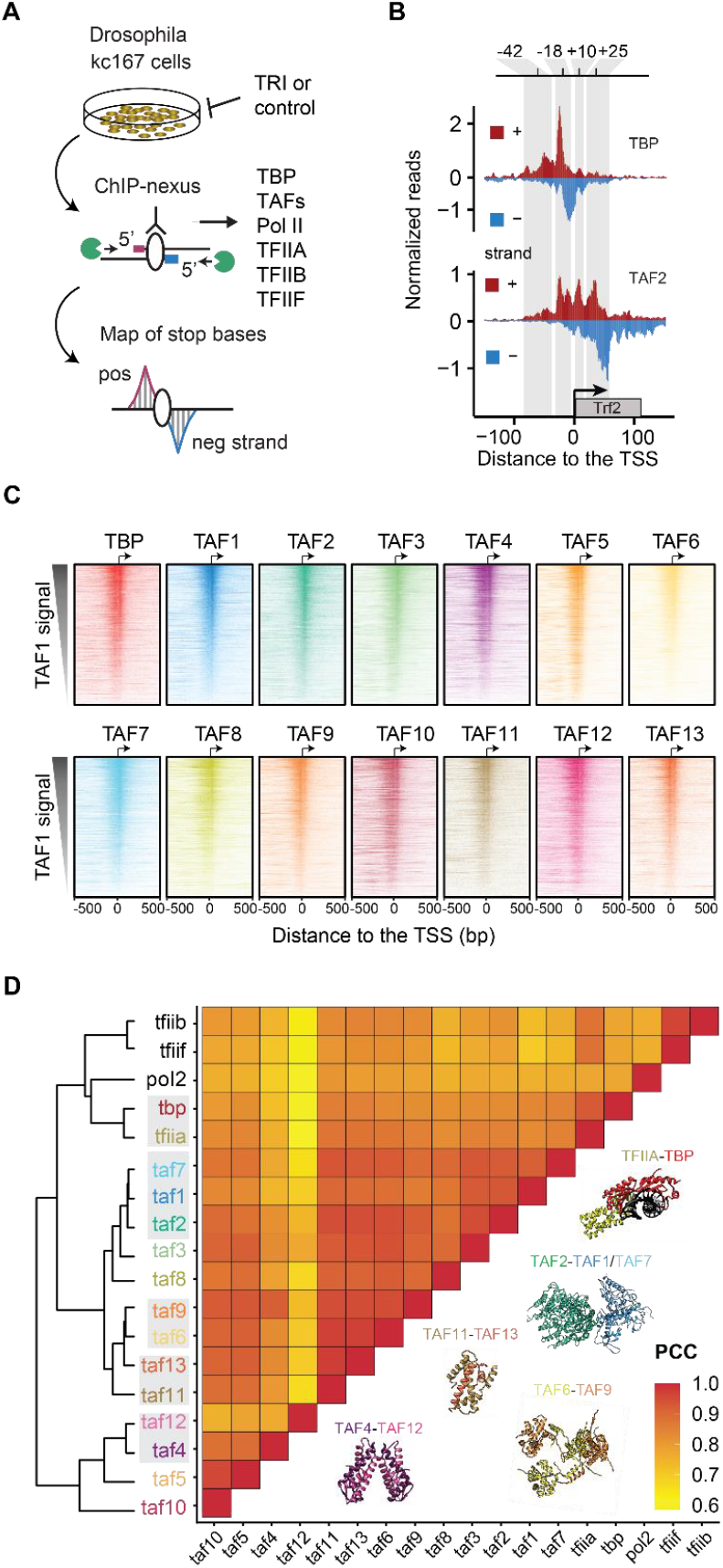
All TFIID subunits are found across active promoters. **A)** Schematic representation of the ChIP-nexus workflow. **B)** TBP and TAF2 binding profile at the *Trf2* gene, shown as normalized reads for each stop base on the positive strand (red) and the negative strand (blue). **C)** Binding of each TFIID subunit across all active promoters with narrow transcription initiation (n=1,769) as determined by CAGE-seq data. Binding to the negative and positive strand is shown in the same color. Promoters were sorted based on the total binding of TAF1 within 100 bp centered on the TSS. **D)** Pearson correlation coefficients (PCC) and hierarchical clustering of the binding of TFIID subunits and other PIC components across all active promoters. The total binding signal was measured within 100 bp centered on the TSS. Note that proteins that physically interact (see structures) often clustered first (marked in grey).

In support of the quality of our data, all experiments showed strong and distinct binding footprints of TFIID subunits at promoter regions (Figure 1B). We then selected all active focused promoters, defined as annotated TSSs where CAGE-seq data in Kc167 cells show transcription initiation in a narrow window (n = 1,769; Figure S3, see Methods). After confirming that the biological replicate experiments were highly reproducible (Figures S1,S2), we analyzed the pooled ChIP-nexus coverage of each TFIID subunit.

When the promoters were sorted by their TAF1 occupancy, the other TFIID subunits followed the same intensity pattern (Figure 1C). TAF9, TAF10 and TAF12 are also components of the SAGA complex, which associates with promoter and enhancer regions^31,32^, but since the ChIP-nexus data are mapped at the TSS and sorted by TAF1 signal, the contributions of SAGA-specific signals from the shared TAFs are likely blurred and minimized.

We next clustered the TFIID subunits based on their binding level similarity across the promoters, measured as pairwise Pearson correlation coefficients (PCCs). We included ChIP-nexus data for the PIC transcription factors TFIIA, TFIIB, TFIIF and PolII for comparison, which showed high correlations with the TFIID subunits (median PCC = 0.9) (Figure 1D). Notably, TBP had a higher correlation with TFIIA and other PIC components than with the TAFs (Figure 1D), consistent with a direct physical interaction between TBP and TFIIA in the PIC^5,6,33,34^.

Importantly, the TFIID subunits were overall all highly correlated in their binding levels (median PCC = 0.9), and the most correlated TAFs (TAF4/12, TAF1/7/2, TAF11/13 and TAF6/9) were known to each form a physical complex^9^ (Figure 1D). This suggests that high correlations are indicative of physical interactions, consistent with formaldehyde fixation generating abundant protein-protein crosslinks^35,36^, causing interacting proteins to be pulled down with the same protein-DNA contacts in ChIP-nexus experiments^30,37–40^. The results indicate that our *in vivo* data align with the known *in vitro* structure of TFIID.

### TFIID structure inferred from *in vivo* footprints

To identify differences by which TFIID subunits contact DNA, we analyzed the base-resolution ChIP-nexus binding footprints. Since interacting proteins are crosslinked to each other and often pulled down together, they produce similar footprints in ChIP-exo/nexus data^30,37–40^. Exact protein-protein interactions cannot be inferred since formaldehyde crosslinks do not occur at all amino acids^36^, but as an accumulative measure, we expect the ChIP-nexus binding profiles to resemble the structural organization of TFIID *in vivo*.

To analyze the ChIP-nexus binding profiles for each TFIID subunit at maximum signal and resolution, we used the ChIP-nexus data of TBP and the 13 TAFs after triptolide treatment and averaged across the promoters with narrow transcription initiation windows (Figure S3). We then identified the main DNA contacts made in these profiles by calculating the midpoint between the peak on the positive strand and the corresponding downstream peak on the negative strand. The main footprint positions were found in six distinct regions, mapped at -150/-50, -30, -18/-14, +10, +19 and +32 bp relative to the TSS (Regions 1-6 in Figures 2A,S4). Moreover, Principal Component Analysis (PCA) grouped the profiles of the 14 TFIID subunits into four modules, which we call the downstream, middle, upstream narrow and upstream broad module (Figures 2A,B). Both the modular organization and the preferred DNA contacts align well with cryo-EM structures of promoter-bound human and yeast TFIID (Figure 2C)^3–6^, demonstrating that our data can provide insights into the structural organization of the complex *in vivo*.

**Figure 2.**
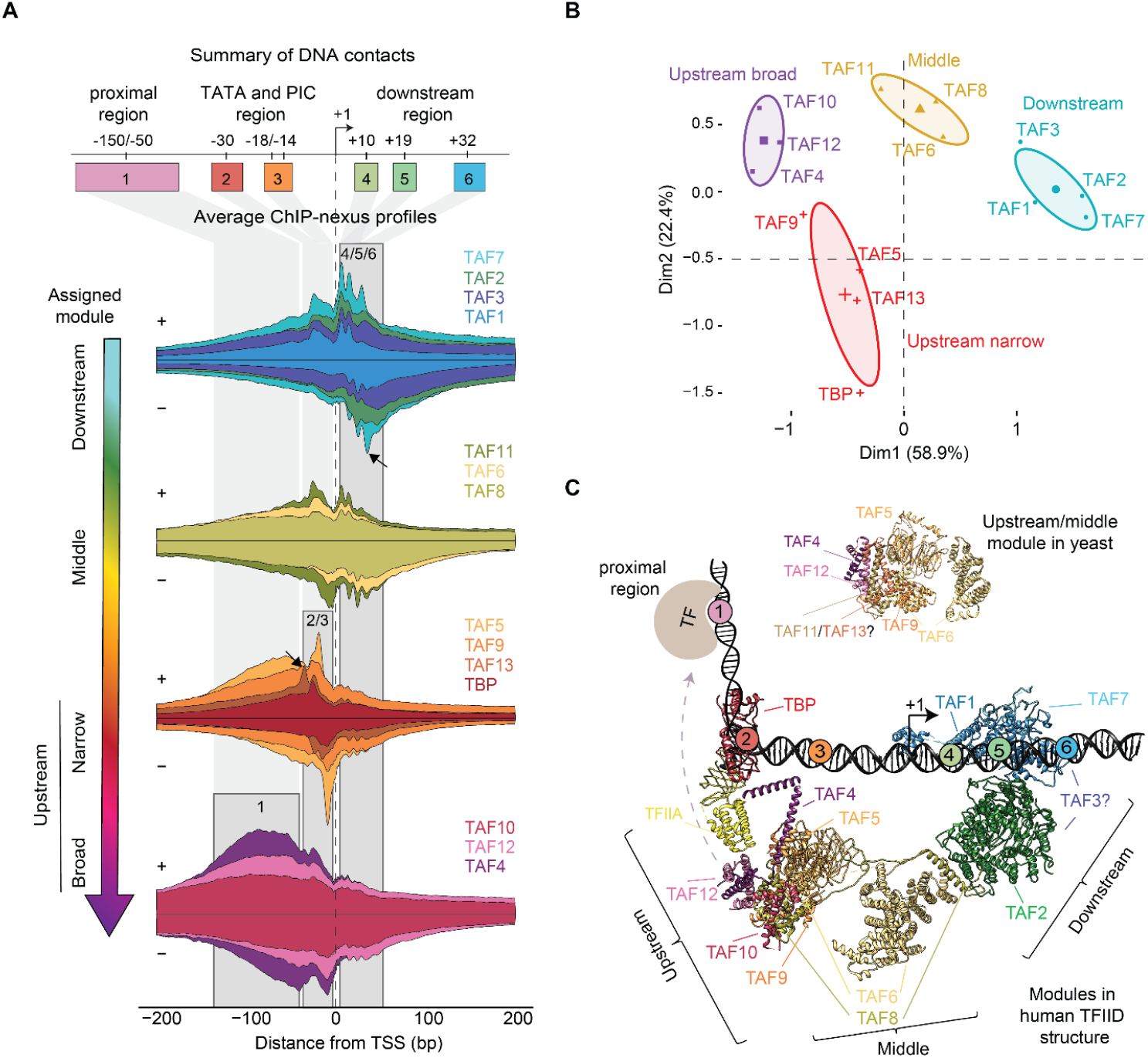
The TFIID ChIP-nexus footprints fall into four modules. **A)** The average ChIP-nexus binding profile of the TFIID subunits centered at the TSS shows a four-module organization of the promoter-bound TFIID complex. For each subunit, the binding profile on the positive strand (top line) and negative strand (bottom line) is shown in the sam e color and stretched to an arbitrary scale to facilitate comparison with other subunits. **B)** Principal Component Analysis (PCA) of the averaged ChIP-nexus profiles confirms the modular organization. **C)** Crystal structure of the yeast (top, lobe B) and human (bottom, lobe B and C) promoter-bound TFIID is consistent with the modular organization and DNA contacts identified by the ChIP-nexus experiments.

Our downstream module is the lobe that engages with DPR core promoter elements (lobe C). TAF1, TAF7 and TAF2 form three clustered footprints at +10, +19, and +32 bp (Figure 2A, regions 4-6), consistent with the human TFIID cryo-EM structures^3,5,6^ and contradicting previous biochemical experiments implicating *Drosophila* TAF6/9 in binding to DPR elements^41^. Interestingly, our downstream module also contains TAF3, which is part of the flexible lobe A and whose location in promoter-bound TFIID is not well established^5,6^. The inclusion of TAF3 together with TAF1/2/7 is supported by crosslinking mass spectrometry analysis in yeast^4^ and consistent with TAF3 containing a PHD finger that interacts with the H3K4me3 mark of the +1 nucleosome downstream of the TSS^42,43^.

The middle module shows footprints both upstream and downstream of the TSS (Figure 2A, regions 2-5). As expected, it contains TAF6 and TAF8, which are known to bridge the upstream and downstream module in cryo-EM structures and thus indirectly contact the promoter both upstream and downstream (Figure 2C). More unexpectedly, TAF11 clusters within the middle module, even though TAF11 forms a heterodimer with TAF13, which is found in the upstream module^44^. TAF11/13 are known to interact with TBP and inhibit its binding to DNA in the canonical state of TFIID. Thus, in the conformation of TFIID where TBP engages the promoter, TAF11/13 must relocate or disengage from TBP^5,6^. Our *in vivo* footprint data are therefore consistent with a rearrangement in which TAF11 becomes positioned closer to TAF6 and TAF8 to allow TBP to bind DNA.

The narrow upstream module consists of TBP, TAF5, TAF9 and TAF13, which are from the two lobes that engage the upstream promoter region (lobe A and B). This module shows strong footprints around the site of PIC formation at -18/-14 bp in our data (Figure 2A, regions 2 and 3). The density for these TAFs is poorer in cryo-EM structures^4–6^, but overall agrees with our data.

Notably, the strongest footprint of TBP is found at the -18 bp position, consistent with previous *Drosophila* TBP ChIP-nexus data and human TBP ChIP-exo data^30,37,38^. This is unexpected since the TATA box is found at -30 bp, and TBP also binds at a similar position at TATA-less promoters in cryo-EM structures^6^. Moreover, TAF13 has its strongest footprint at the -30 bp position (Figure 2C, arrow), suggesting that the underlying sequence is conducive to ChIP-nexus footprints. This suggests that TBP crosslinks poorly at the TATA box, likely because TBP primarily interacts with the DNA phosphate backbone^45,46^, rather than the bases, which are the preferred substrate of formaldehyde crosslinking^36,47^. TBP’s crosslinking signal at -18 bp likely originates from TBP’s interaction with PIC components like TFIIB, which contact DNA downstream of the TATA box^6,45,48^. We will examine this TBP footprint in more detail later.

Finally, we identified a broad upstream module, which shows high levels of diffuse footprints upstream of the TSS (Figure 2A, region 1). It consists of TAF4, TAF10, and TAF12, which are present in two copies and belong to the two lobes that engage the upstream promoter region^3,5,6^. These diffuse upstream footprints might be due to contacts with enhancers upstream of the TSS since many TAFs are known to engage with DNA-bound transcription factors^49–58^. Additionally, TAF10 and TAF12 could also engage these regions as part of the SAGA complex^59^.

Taken together, the ChIP-nexus profiles show strong agreement with the in vitro cryo-EM structures, provide insights for less well-characterized subunits such as TAF3, and point to unexplained observations for TAF11/13 and TBP.

### The TBP binding profile is promoter-type-specific

The high correlation of the TFIID subunits across promoter types argues against partial TFIID complexes at specific promoters but does not rule out subtle differences by which TFIID subunits, especially TBP, interact with different promoter types. To test this, we analyzed our ChIP-nexus data from untreated cells to capture all possible promoter states (see Figure S6 for data from triptolide-treated cells). We identified four representative promoter types among our defined set of narrow promoters, selected based on the presence of core promoter elements at specific positions relative to the TSS, as done previously^30,60,61^ (Figures 3A, S5).

**Figure 3.**
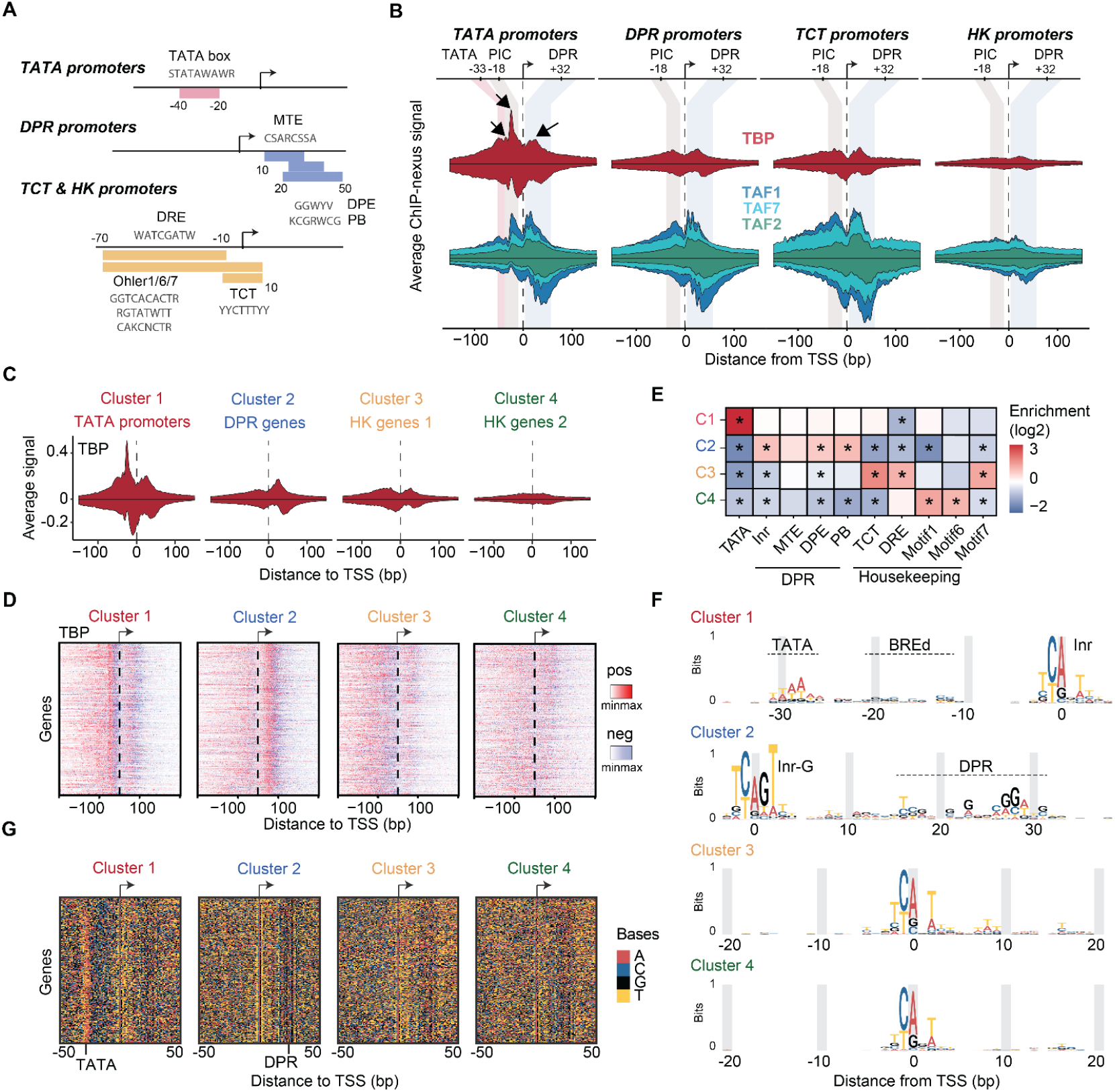
TBP is promoter-specific and allows the *de novo* identification of promoter types. **A)** Schematic summarizing the different promoter elements used for analysis (see Methods table 1 for more information). **B)** The average ChIP-nexus binding profile of TBP (top; red), TAF1, TAF2 and TAF7 (second row) at TATA, DPR, TCT and HK promoters. **C)** Average ChIP-nexus binding profiles of TBP at the four different k-means clusters. **D)** Distribution of TBP binding location at all promoters within each promoter cluster. Binding to the positive and negative strand is shown in red and blue, respectively. Signal per TSS was normalized to cluster promoters based on the TBP profile shape (see Methods). **E)** Promoter clusters are enriched for different core promoter elements. Statistical significance was calculated using a Fisher’s exact test with multiple-testing correction, ^*^*p* < 0.05. **F)** Position weight matrix (PWM) of the identified upstream, initiator and downstream regions of the four identified promoter clusters. **G)** Underlying sequence information contents across the promoter clusters colored by base.

TATA promoters (n = 191) have a TATA box located -30 bp upstream of the TSS. Downstream promoter region (DPR) promoters (n = 334) have a DPR element at +20-30 bp, thus an MTE, DPE or PB motif. DPR promoters typically show higher levels of Pol II pausing^30,61,62^. TCT promoters (n = 78) initiate at a TCT motif and are often found at ribosomal genes^63,64^. Finally, housekeeping (HK) promoters (n = 282) have at least one housekeeping core promoter element (DRE, Ohler 1, 6 and 7 motifs) and no TATA, TCT or DPR element. HK promoters tend to be more dispersed with broader initiation windows^65–68^, but this dispersion was kept minimal by our selection of promoters with narrow transcription initiation.

Comparing the profiles of all TFIID subunits at the four promoter types revealed that TBP has the most promoter-specific profile (Figure 3B), although there are subtle differences among the TAFs, e.g., the binding at the TCT and HK promoters is more diffuse. The fact that TBP stands out as being most promoter-specific supports the hypothesis that TBP is a mobile component of TFIID^1,9,69^ and may indicate functional differences between promoter types.

Notably, the strong footprint of TBP at the upstream -18 bp position is specific for TATA promoters (Figure 3B). It is strongest after triptolide treatment, supporting its association with the PIC (Figure S6). TBP also displays a broad footprint +32 bp downstream of the TSS (Figure 3B). This footprint is present at all promoter types to various degrees, but is not observed after triptolide treatment (Figures 2A, S6, S7). This suggests that downstream TBP does not occur during PIC assembly but during another step of the transcription cycle, though we can exclude its association with stable Pol II pausing (Figure S9). We note that cryo-EM studies have suggested that TBP is in a promoter-downstream position before it gets loaded onto the upstream position^3,5,6^, thus it is possible that downstream TBP represents the TFIID loading state.

While TBP appears to have the most promoter-specific profile, this conclusion is so far based on average profiles across selected promoter sets. To test whether the differential TBP binding profile is robust across individual promoters, we asked whether we could re-classify promoters based on their individual TBP binding profile. Using the base-resolution TBP profiles at the previously defined narrow promoters (Figure S3), we performed k-means clustering allowing four possible clusters (see Methods).

The TBP profiles of the four clusters (Figures 3C,D) and their enrichment of core promoter elements (Figure 3E) suggest that the clusters represent TATA promoters (cluster 1), DPR promoters (cluster 2), and two sets of TCT/HK promoters, one with diffuse TBP ChIP-nexus footprints (cluster 3), and the other with low TBP ChIP-nexus footprints (cluster 4) (Figures 3C,D).

As further validation, we asked whether we could *de novo* discover the core promoter elements from each of the promoter sets clustered by TBP profile (Figure 3F,G). We aligned the sequences at the TSS and visualized the bases coded by color. Due to their strict positioning relative to the TSS, certain core promoter elements were discernible by eye (Figure 3G). Clusters 3 and 4 (TCT/HK) showed clear Inr elements, while clusters 1 and 2 showed additional strong sequence patterns indicative of TATA and DPR promoters, respectively (Figure 3G).

The resulting motif logos also closely matched the known core promoter elements^70–72^ (Figure 3F). Notably, the TATA and DPR promoters both have Inr elements, but only those from DPR promoters were enriched for the G at position +2 (Inr-G) as previously reported^61^. Furthermore, a CG-rich sequence next to the TATA motif resembles the TFIIB response element (BREd)^48^, but we cannot exclude that this sequence simply enhances the crosslinking of TFIIB at this position. Taken together, we conclude that TBP displays binding profiles that can be used to classify promoter types *de novo* and predict their sequence elements.

### TAF-independent PIC formation at TATA promoters

We next investigated what causes the strong upstream footprint of TBP at TATA promoters. We considered that it might be due to prolonged binding of TBP at the TATA box, which could explain the more rapid re-initiation at TATA promoters^73–77^. If so, one would expect higher total levels of TBP relative to the transcriptional output at TATA promoters. To test this, we normalized the ChIP-nexus binding profiles of TBP and the other factors by dividing the binding signal at each promoter by the corresponding CAGE-seq transcript level.

Once normalized, the levels of TFIIA and TFIIB were not statistically different between TATA and DPR promoters (Figures 4A,B), suggesting that PIC factors occupancy is roughly proportional to the amount of transcripts initiated at these promoters. An exception were TCT promoters, which had lower levels of TFIIA and higher levels of TFIIB, indicating functional differences (Figures 4B). TRF2 was also significantly lower at TATA promoters, consistent with TRF2 not binding the TATA box and with previous ChIP experiments^32,63,64,67,78,79^. Importantly, TBP followed the trends of the PIC factors and was not significantly higher at TATA *versus* DPR promoters after normalization (Figure 4B), which supports the idea that TBP does not remain bound through multiple initiation events at TATA promoters.

**Figure 4.**
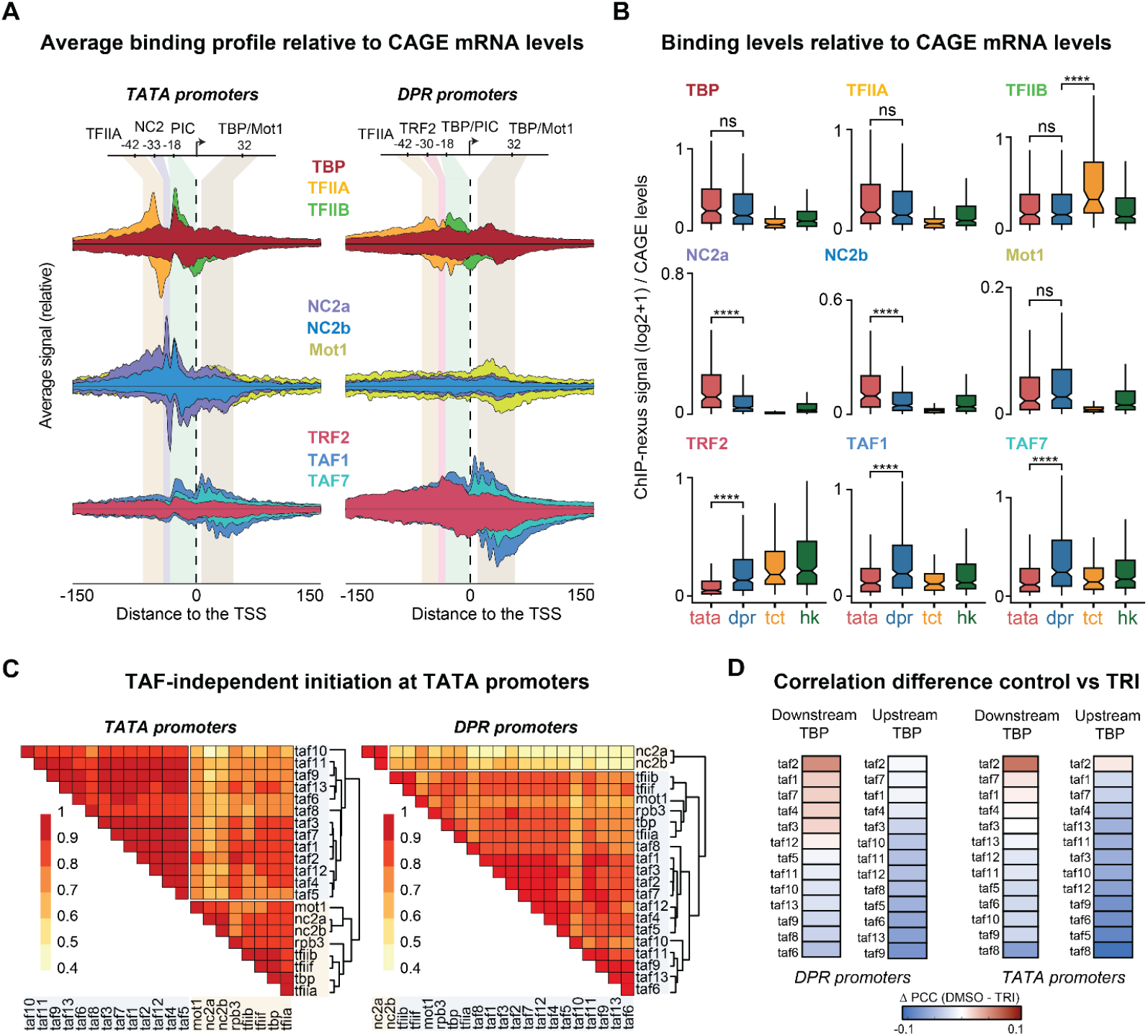
TFIID-independent initiation at TATA promoters. **A)** Average transcription-normalized binding profile of TFIIA, TFIIB, (top row; yellow, grey and green), TBP, NC2a, NC2b and Mot1 (middle row; red, dark blue, purple and pink), TAF1, TAF7 and TRF2 (third row; blue, turquoise and orange) at TATA and DPR promoters. Transcrip tion normalization was calculated by dividing the bp signal at each promoter by its corresponding CAGE-seq score. **B)** Transcription-normalized signal distribution for the transcription factors shown in panel A (Wilcoxon test). Occupancy was calculated dividing the total ChIP-nexus signal (calculated 100 bp around the TSS) at each promoter by the corresponding CAGE-seq levels. **C)** Pearson correlation heatmaps plots for PIC components, NC2 and Mot1 at TATA and DPR promoters. **D)** Heatmaps showing the difference in Pearson correlation coefficients (PCC) between control (DMSO) and triptolide (TRI) when correlating upstream or downstream TBP with TAFs. Results are shown for DPR and TATA *promoters*.

We next investigated whether TBP-associated proteins might provide clues to TBP’s promoter-specific distribution. We performed ChIP-nexus experiments on Mot1 and both NC2 subunits, two factors known to alter TBP binding on DNA^20,21,23,80,81^ (Figures 4C, S8A). Mot1 showed elevated binding at all locations where TBP binds at TATA and DPR promoters, especially at the downstream footprint (Figures 4A; S8A). In contrast, both NC2 subunits, NC2α and NC2β, were specifically bound to TATA promoters, significantly higher than at DPR promoters even after normalization (p < 0.001) (Figures 4A,B). Each NC2 subunit showed two sharp ChIP-nexus footprints: one at -18 bp, coinciding with that of TBP, and another at -33 bp near the TATA box (Figure 4A; S8A). These results match remarkably well the crystal structure of TBP-NC2-DNA ternary complex (Figure S8B)^82^. This could be because NC2α and NC2β contain lysine residues positioned to cross-link with DNA in the vicinity of the TATA box: NC2α contacts the TATA box underneath TBP (Lys18, Lys19), whereas NC2β also interacts with the downstream region (Lys63, Lys64)^82^. This likely explains why NC2 generates ChIP-nexus footprints at the TATA box where TBP cannot.

Finally, we investigated whether the strong upstream TBP footprint at TATA promoters could represent TBP as part of a TAF-independent PIC. This is in agreement with classic studies showing that TATA promoters do not require TAFs for transcription initiation *in vitro^83–86^*, as well as with studies showing that TATA promoters have reduced TAF occupancy and a lower dependency on TAFs for transcription^13,37,84,87,88^. When we analyzed the transcript-normalized profiles of the TAFs, we found that all TAFs showed significantly lower levels at TATA promoters. Since TBP, TFIIA and TFIIB were nevertheless present at similar levels at TATA and DPR promoters (Figures 4A,B and S7), these results support the hypothesis that TBP can form a PIC without TFIID on TATA promoters.

To obtain additional evidence not based on binding differences between promoter types, we analyzed each promoter type separately and performed a pairwise correlation analysis between the binding levels of TBP, TAFs, TFIIA, TFIIB, TFIIF, NC2 and Mot1 (Figure 4D). At TATA promoters, TBP shows stronger correlations with TFIIA, TFIIB, and TFIIF, NC2 and Mot1 than with the TAFs, implying that TBP does not always co-occur with the TAFs (Figure 4D, left). This pattern was not observed at DPR promoters, where the binding levels of all factors were more homogenous, consistent with TFIID being an integral part mediating PIC formation at DPR promoters (Figure 4D, right).

Taken together, these results suggest that the promoter-specific profile of TBP can be explained by TBP’s ability to mediate PIC formation at TATA promoters by directly binding the TATA box rather than being loaded by TFIID from the downstream to the upstream position of the promoter. Since TAFs are nevertheless present, these results suggest that TATA promoters can mediate TAF-dependent and TAF-independent PIC formation *in vivo*.

Interestingly, this conclusion suggests that downstream TBP is associated with TFIID. TATA promoters not only have lower levels of TAFs but also lower levels of downstream TBP compared to its upstream binding. In contrast, DPR promoters, which have downstream sequences that favor TFIID binding, have the highest levels of downstream TBP. This supports our hypothesis that downstream TBP is part of the TFIID loading state.

If downstream TBP represents a TFIID loading state, we wondered whether we could identify TAFs that correlate more highly with the TFIID loading state and thus might be physically close to TBP when downstream. For this, we took advantage of the fact that TBP is found downstream under steady-state control conditions, but not after triptolide treatment. We asked which TAFs have the highest increase in correlation with downstream TBP under control *versus* triptolide conditions.

We found TAF2 to have the greatest increase in correlation, followed by TAF1 and TAF7, and was overall higher at DPR promoters versus TATA promoters (Figure 4D), consistent with the downstream module mediating the loading of TBP. We would have expected TAF1 to have the highest signal since it can contact TBP directly through its TAND domain, but it is possible that this is because the Lys-Arg-enriched DNA-facing surface of TAF2 is more easily cross-linked to DNA than the DNA-binding surface of TAF1.

These findings propose a model that explains the strong upstream TBP footprint at TATA promoters and the higher downstream footprint at DPR promoters. At both promoter types, TFIID initiates transcription in a similar way, by relocating TBP from the downstream to the upstream promoter position (Figure 5). At TATA-containing promoters, however, TBP can directly bind to the TATA box and mediate PIC formation without the TAFs. This mode of TFIID-independent initiation produces sharper footprints because TBP is more precisely positioned at the TATA box, while also reducing the downstream TBP binding that occurs when TBP is loaded by the TAFs.

**Figure 5.**
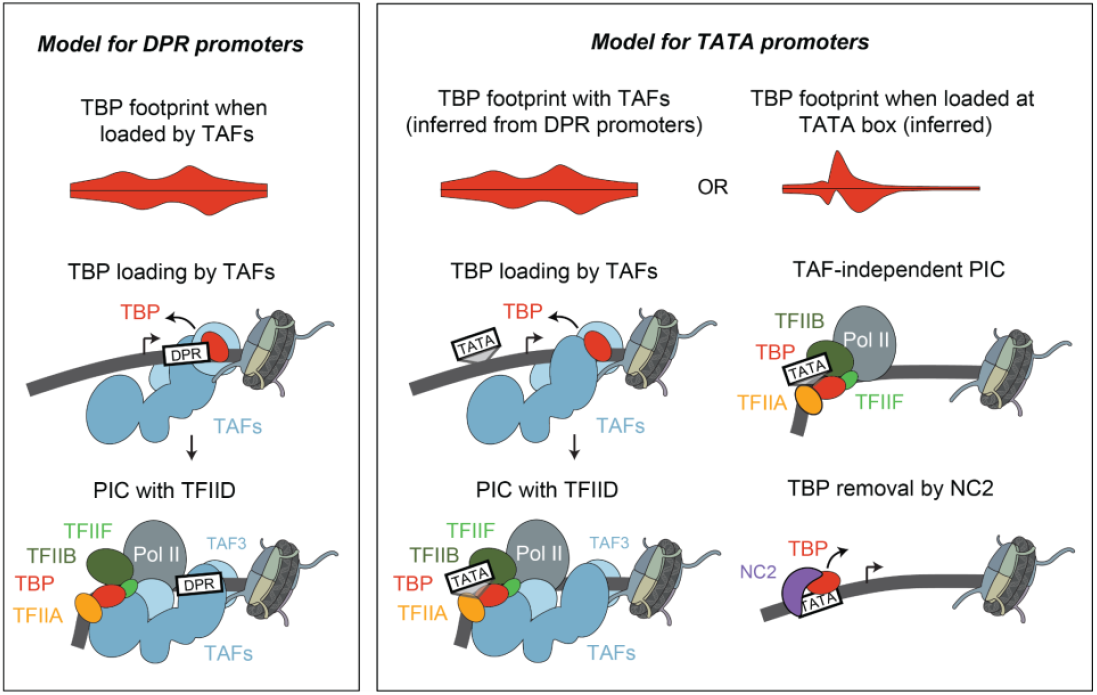
Model for transcription initiation at TATA and DPR promoters.

## Discussion

While cryo-EM studies have unraveled the structure of TFIID bound to specific promoters in yeast and humans, it has remained speculative how TFIID’s structure and function might differ between promoter types. By mapping all 14 TFIID subunits at base-pair resolution, we provide the first comprehensive view of how TFIID binds across different promoter types *in vivo*. We found the data remarkably consistent with cryo-EM structures, suggesting that the holo-TFIID complex is largely found at active promoters, in line with recent functional and structural investigations^6,89–91^. We found however that TBP displays significant promoter-specific differences, allowing us to connect structural studies with previous observations from biochemistry and genetics and construct a model of the function of TFIID at different promoter types.

Our *in vivo* ChIP-nexus profiles support a dual mechanism of initiation at TATA promoters, one that is TAF-dependent and one that is TAF-independent. By normalizing the signal of TFIID and PIC components at each promoter to its transcription output, we found that TATA promoters have significantly lower TAF occupancy relative to their transcriptional activity, consistent with a substantial fraction of their transcription being driven in a TAF-independent manner. Furthermore, we find that the TBP high-resolution binding profile at TATA promoters matches two modes of initiation. On one hand, a strong TBP upstream footprint is specific to TATA promoters and represents TAF-independent initiation. On the other hand, all promoters, including TATA promoters, have a downstream TBP footprint during steady-state conditions, which likely represents the DNA-bound loading state of TFIID that allows TBP to be placed to the upstream position^5^. The combination of these two initiation modes largely explains the promoter-specific binding of TBP (Figure 5).

By obtaining *in vivo* evidence of both TAF-dependent and TAF-independent transcription at TATA promoters, we provide clarity to a long-standing debate of TFIID in the field. It has been debated whether TFIID is essential for transcription initiation and how it depends on the promoter type^84,85,92–94^. There has been evidence in yeast for TAF-dependent and TAF-independent forms of TBP on promoters^95,96^, but early genome-wide experiments suggested that removing TAFs reduces transcription at both TATA and TATA-less genes, though a subset of TATA-rich promoters were shown to be less dependent on TAFs^90,97^. Subsequent work in yeast and mammals further consolidated that transcription of many TATA-containing genes persists upon depletion of certain TAF subunits^42,87,96^. Similarly, more recent genome-wide analyses found promoters where TBP and Pol II remained bound or active even when a TAF showed minimal occupancy^37,42,87,88,96,98^. Thus, our results are consistent with these studies and provide the resolution necessary to directly separate TAF-dependent and TAF-independent initiation at TATA promoters, along with providing *in vivo* evidence for a TFIID loading state.

Our high-resolution profiles also provide more conclusive binding information on NC2 and the TBP-related factor TRF2. We detected strong and sharp footprints for both NC2 subunits at TATA promoters, but not at TATA-less promoters. Such preference for TATA was not observed with lower resolution ChIP data in yeast^21^, but it is consistent with newer yeast ChIP-exo data^39^. TRF2 showed the opposite behavior; its binding was particularly low at TATA promoters and higher at DPR, TCT and HK promoters, in agreement with previous studies^32,63,78,99–101^.

For NC2, our data support a model where it predominantly recognizes the TATA-bound TBP complex to suppress excessive initiation at TATA promoters. This is consistent with evidence that NC2 prevents PIC assembly and destabilizes TBP binding at the TATA box^21,82^. From this perspective, observations of NC2 affecting TATA-less promoters^21–23,102,103^ could be indirect: disruption of NC2 could increase TBP’s usage at TATA promoters and sequester it away from TATA-less promoters. However, it is also plausible that NC2 (together with Mot1) has a broader role in removing TBP from nonspecific sites^104–106^ since such transient, non-specific binding of TBP, NC2 and Mot1 would not cause specific footprints.

Finally, our findings offer an opportunity to consider models of promoter-specific behaviors. TATA-containing promoters often drive high bursts of transcription, whereas TATA-less promoters typically lead to steadier outputs^12,60,107–111^. It has been proposed that TBP’s long residence time at TATA promoters enables multiple rounds of initiation, yielding large bursts^73–77^. However, recent structural analyses show that TBP, together with TFIIA, TFIIB and TFIIF, is evicted from DNA as Pol II transitions into elongation^112^. Similarly, *invivo* experiments in yeast demonstrate that inactive pre-initiation complexes rapidly dissociate genome-wide when transcription is stalled^87^. These findings imply that TBP must rebind *de novo* for each round of transcription during a burst.

In light of these findings, our results provide potential explanations for the larger transcriptional bursts observed at TATA promoters. By virtue of having both TAF-dependent and TAF-independent options, TATA promoters may exhibit more rapid reloading of TBP during a transcription burst. Once TBP is evicted by an elongating polymerase, the promoter can quickly reacquire TBP either through TFIID (which might remain nearby and redeploy TBP)^10,113^ or via direct TBP binding to the TATA box. While having two mechanisms should promote robust re-initiation^113^, it is possible that the higher burst amplitude is mainly driven by TAF-independent re-initiation, which is faster^114^. Depletion of TAF1 in *Drosophila* cells leads to reduced transcription at DPR promoters and to larger and prolonged transcriptional bursts at TATA promoters^115^. While we cannot exclude indirect effects, i.e. that reduced transcription upon TAF1 depletion makes more PIC components available for TATA promoters, the results could indicate that TAF-dependent initiation constrains the upper limit of the burst amplitude. This is consistent with TFIID mediating a steadier transcription output at DPR promoters.

In summary, our high-resolution binding data of TFIID provide new details that can be used to revisit and reininterpret previous experimental results, leading to revised or new models for the function of promoters.

## Methods

### Cell culture and transcription inhibitor treatment

Drosophila melanogaster Kc167 cells (purchased from DGRC; negative for mycoplasma contamination) were grown at 25 °C in HyClone SFX-Insect Cell Culture Media. Transcription inhibitors were added directly into SFX media. Cells were treated with 10 μM triptolide (TOCRIS Bioscience, 3253; dissolved in DMSO) at room temperature for 1h. Equivalent amounts of DMSO treatment (2%) were used as the control.

### ChIP-nexus experiments

For each ChIP-nexus experiment, 10^7^ Kc167 cells were fixed with 1% formaldehyde in culture media at room temperature for 10 min with rotation. Fixed cells were washed with cold PBS, incubated with Orlando and Paro’s Buffer (0.25% Triton X-100, 10 mM EDTA, 0.5 mM EGTA, 10 mM Tris-HCl pH 8.0, with freshly added Protease Inhibitor) for 10 min at room temperature with rotation, and then centrifuged and re-suspended in ChIP Buffer (10 mM Tris-HCl, pH 8.0; 140 mM NaCl; 0.1% SDS; 0.1% sodium deoxycholate; 0.5% sarkosyl; 1% Triton X-100, with freshly added Protease Inhibitor). Sonication was performed with a Bioruptor Pico for five rounds of 30 s on and 30 s off. Chromatin extracts were then centrifuged at 16,000 g for 5 min at 4° C, and supernatants were used for ChIP.

To couple Dynabeads with antibodies, 50 µl Protein A and 50 µl Protein G Dynabeads were used for each ChIP-nexus experiment and washed twice with ChIP Buffer. After removing all the liquid, Dynabeads were resuspended in 400 µl ChIP Buffer. 10 µg antibodies were added, and tubes were incubated at 4°C for 2 hr with rotation. After the incubation, antibody-bound beads were washed twice with ChIP Buffer.

For chromatin immunoprecipitation, chromatin extracts were added to the antibody-bound beads and incubated at 4°C overnight with rotation and then washed with Nexus washing buffer A to D (wash buffer A: 10 mM Tris, 1 mM EDTA, 0.1% Triton X-100, wash buffer B: 150 mM NaCl, 20 mM Tris-HCl, pH 8.0, 5 mM EDTA, 5.2% sucrose, 1.0% Triton X-100, 0.2% SDS, wash buffer C: 250 mM NaCl, 5 mM Tris-HCl, pH 8.0, 25 mM HEPES, 0.5% Triton X-100, 0.05% sodium deoxycholate, 0.5 mM EDTA, wash buffer D: 250 mM LiCl, 0.5% IGEPALCA-630, 10 mM Tris-HCl, pH 8.0, 0.5% sodium deoxycholate, 10 mM EDTA). End repair and dA-tailing were performed using the NEBNext End Repair Module and the NEBNext dA-Tailing Module. ChIP-nexus adaptors with mixed fixed barcodes (CTGA, TGAC, GACT, ACTG) were ligated with Quick T4 DNA ligase and converted to blunt ends with Klenow fragment and T4 DNA polymerase. The samples were treated with lambda exonuclease and RecJf exonuclease. After each enzymatic reaction, the chromatin was washed with the Nexus washing buffers A to D and Tris buffer (10 mM Tris, pH 7.5, 8.0, or 9.5, depending on the next enzymatic step). The chromatin was then eluted and subjected to reverse crosslinking and ethanol precipitation. Purified single-stranded DNA was then circularized with CircLigase, annealed with oligonucleotides complementary to the BamHI restriction site and linearized by BamHI digestion. The linearized single-stranded DNA was purified by ethanol precipitation and subjected to PCR amplification with NEBNext High-Fidelity 2X PCR Master Mix and ChIP-nexus primers. The ChIP-nexus libraries were then gel-purified before sequencing with Illumina NextSeq 500.

### ChIP-nexus data processing

ChIP-nexus data generated in this work and reprocessed from GSE85741^30^ were first preprocessed by trimming and reassigning fixed and random barcode sequence information, stored as read names for use in downstream processing. Reads were next trimmed to remove adapter sequences using Cutadapt (v.4.2)^116^ and aligned to the dm6 genome assembly using bowtie2 (v.2.3.5.1)^117^. Using aligned coordinates alongside the stored fixed and random barcode sequence information, reads were deduplicated. Coverage was generated by collecting only the first “stop” base of each aligned read. In order to compare reproducible, base-resolution footprinting profiles, each replicate ChIP-nexus experiment was first normalized via reads-per-million (RPM) scaling of coverage to the total aligned reads of a replicate experiment. Next, normalized replicate footprints were pooled and averaged by number of replicates per TF and condition.

### CAGE-seq data processing and Drosophila TSS annotation

Publicly available CAGE-seq data from *Drosophila* melanogaster Kc167 cells (modENCODE: ENCSR827COJ) were processed using the STAR aligner (v2.7.1a)^118^ against the *Drosophila* melanogaster UCSC dm6 reference genome. The resulting coordinate-sorted BAM files were analyzed using the *CAGEr* (v2.14.0) package in R to obtain TSSs (Transcription Start Sites)^119^. Quality control was assessed using Pearson correlations and biological replicates were merged and normalized using a power-law approach (*α = 1*.*19, T = 10*^*6*^) within the fit range of 3–40,000 counts.

Individual replicate TSSs were grouped into tag clusters within each sample using distance-based clustering (*distclu*, CAGEr package) with a maximum merging distance of 30 bp. To delineate precise coordinates for each tag cluster, the cumulative TSS signal was calculated and the 10th and 90th percentiles of the cumulative distribution were determined to delineate the central coordinates which contain 80% of transcriptional signal.

Tag clusters with a TPM ≥ 0.5 in at least one sample were then merged across replicates (*aggregateTagClusters*, CAGEr package) with a maximum inter-cluster distance of 100 bp to define consensus TSS clusters. The genomic coordinates of each consensus cluster were defined as the union of the constituent clusters’ central activity regions. Finally, cumulative TSS distributions and quantile positions were recomputed for the consensus clusters to refine their boundaries. The dominant TSS of each consensus cluster was identified as the genomic position with the highest summed TSS signal across the two replicate samples. For promoters that exhibited a single TSS, only those with ≥3 TPM were kept.

TSSs from FlyBase (r6.64, FB2025_03) were used to annotate the TSS clusters. If multiple clusters mapped to the same gene or transcript, only the cluster with the highest TPM was selected. In total, we identified 6,314 active promoters. Active focused promoters where transcription initiation occurred within a narrow window (n = 1,769) used for subsequent analysis were selected based on their interquantile cluster width (< 11 bp).

### RNA-seq library preparation

Total mRNA were extracted for Drosophila melanogaster Kc167 cells using Invitrogen TRIzol reagent. cDNA libraries are prepared using Illumina TruSeq Stranded mRNA Preparation Kit with polyA enrichment and sequenced on Illumina NextSeq 500.

### RNA-seq data processing

All RNA-seq pair-ended sequencing data were aligned to the UCSC dm6 reference genome using STAR (2.7.1a)^118^ and raw read counts per gene were quantified with “--quantMode GeneCounts” option. RNA-seq levels are TPM-normalized across replicates, measured using the UCSC dm6 transcriptome annotations^120^.

### Principal Component Analysis

Average ChIP-nexus binding profiles under triptolide treatment were quantified per base pair within a 101 bp window centered on each annotated TSS. The average signal for each ChIP-nexus binding signal (*x*) was normalized via *x* − *minx*) / *max*(*x*) − *minx*) and subjected to principal component analysis (PCA) using the function *prcomp (center = TRUE, scale = FALSE)* from the base stats package in R.

### Promoter type characterization

Promoter types were selected based on the presence or absence of the motifs listed in Table 1. Each genomic sequence had to match the consensus sequence, with up to one mismatch for the TATA box and zero mismatches for the other elements. This sequence matching occurred at each promoter within the coordinate boundaries reported in Table 1. TATA and TCT promoters were only required to contain the TATA box or the TCT, respectively, and were not excluded for having other elements. DPR promoters were allowed the presence of MTE, DPE and PB, but not TATA, TCT, Ohler1/6/7, or DRE. Housekeeping promoters were filtered to not have motif elements from any other of the promoter categories. Thus, DPR promoters were allowed the presence of MTE, DPE, and PB, but not TATA, TCT, Ohler1/6/7, or DRE, whereas housekeeping promoters were allowed to contain Ohler1/6/7 or DRE, but not TATA, MTE, DPE, PB or TCT.

**Table 1.**
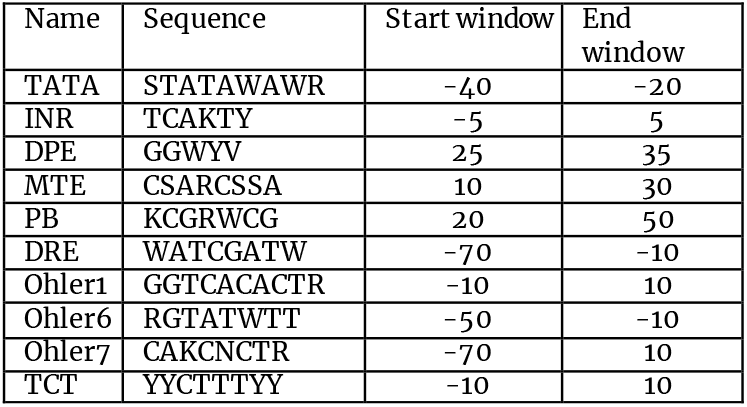

### TBP profile clustering

To cluster promoters based on the TBP ChIP-nexus profile we selected the top 90% narrow promoters given the total TBP ChIP-nexus signal across an 81 bp window centered at the TSS.

To emphasize binding pattern shape, signals for each promoter and position relative to the TSS were ranked from the smallest to the largest values and divided by the number of columns, so the transformed values went from 0 to 1. We then performed K-means clustering (*centers=4, nstart=100 and x=30*) using the base stats package in R.

### Promoter element enrichment

Known *Drosophila* promoter elements in each promoter were identified by the presence of the known consensus sequence with zero mismatches in a specified window relative to the TSS (Table 1). For each k-means cluster and each promoter element, the enrichment was calculated by determining the ratio between the fraction of promoter elements in the cluster and the fraction of the same promoter element in the other four clusters. The significance for the observed frequencies was calculated with Fisher’s exact test and corrected for multiple testing with the Benjamini–Hochberg method.

### Normalization of the ChIP-nexus binding by CAGE-seq expression

To visualize average binding effects around active TSSs, the ChIP-nexus signal was scaled relative to the measured CAGE-seq expression score across the corresponding promoter (−150 bp to +150 bp). Next, ChIP-nexus signal for each promoter was measured around the TSS (-50 bp to +50 bp) and scaled to the promoter’s CAGE-seq expression score in order to compare signals between promoter types. Outliers were excluded from the plot for visualization.

### Hierarchical clustering of TFIID and PIC binding

To compare similarity of ChIP-nexus binding of TFIID subunits and PIC components, binding levels were measured around annotated TSSs (-50 bp to +50 bp) and pairwise Pearson correlations (log2 + 1). Correlations were next transformed to distance values (1−r) and hierarchically clustered using the Ward’s D linkage method. Hierarchical clustering of binding correlations were performed across all narrow promoters (Figure 1D), as well as for each promoter type separately (Figure 4C).

### Paused Pol II half-life measurement

Paused Pol II half-life was re-measured at CTSS promoters with narrow transcription initiation windows from Kc167 cells^30^ with an updated genome (UCSC dm6). Briefly, to analyze the half-life of paused Pol II from these data, promoters were selected if **1)** the total Pol II signal was high (average ChIP-nexus signal >20), **2)** a typical footprint was observed for Pol II (distance between positive- and negative-strand maximum < 80 bp) and **3)** the position of the Pol II footprint was less than 80 bp downstream of the TSS. Upon re-processing, 1,307 promoters fulfilled these criteria. For each promoter, the Pol II ChIP-nexus signal was calculated in a 51 bp window centered on the pausing position (the midpoint between Pol II–positive and Pol II–negative maximum). To calculate the half-life of paused Pol II at each promoter, previously generated Pol II time-course measurements^30^ were fitted into an exponential decay model using nonlinear regression. Promoters with a paused Pol II half-life of longer than 60 min were floored to 60 min to eliminate inflated values due to noise. The 1,307 promoters were then ranked based on half-life and divided into five quintiles. Promoters within quintile 1 (n = 262; Pol II pausing half-lives < 6min) were selected as promoters with short Pol II pausing and promoters within quintile 5 (n = 262; Pol II pausing half-lives > 45min) were selected as promoters with long or stable Pol II pausing.

### Public data sets used for analysis

All protein structures, the DNA-bound human TFIID (pdb_00006mzm), yeast TFIID (pdb_00006hqa), TFIIA-TBP (pdb_00001nvp), TAF1-7-2 (pdb_00006mzm), TAF4-12 (pdb_00001h3o), TAF6-9 (pdb_00006mzd), TAF11-13 (pdb_00001bh8) and NC2-TBP-DNA (pdb_00001jfi), were downloaded from the RCSB Protein Data Bank and colored appropriately using UCSF CHIMERA^121^. CAGE-seq data were downloaded from modENCODE (ENCSR827COJ). Previously published TBP, Poll II and Nelf ChIP-nexus are available under GEO accession GSE85741.

## Data availability

The raw RNA-seq and ChIP-nexus data published in this study are available under GEO accession numbers GSE314461. Gene expression table counts and ChIP-nexus tracks are readily available under the same GEO accession. The analysis code is available on GitHub at https://github.com/zeitlingerlab/G-M_Alcantara_TFIID_2025.

## Acknowledgments

We thank Wanqing Shao for initial assistance with data generation and analysis, and James Kadonaga for providing the TBP, NC2, and Mot1 antibodies. We are also grateful to Marc Timmers, Joan and Ron Conaway, Ralf Jansen, and Ferenc Mueller for their valuable feedback on the manuscript.

## Author Contributions

S.G-M.A. and J.Z. conceived the project. S.G-M.A. performed all the experiments and data analysis. S.B. and M.W. implemented the analysis pipeline and reprocessed the data. J.Z. supervised the project. S.G-M.A., S.B., M.W. and J.Z. prepared the manuscript.

## Funding

This work was funded by the Stowers Institute for Medical Research (SIMR), and supported by NIH grant no. R01HG010211 to J.Z.

## Declaration of interest

J.Z. owns a patent on ChIP-nexus (US-10287628). All other authors declare no competing interests.

