## Supplementary figures for "High-resolution binding data of TFIID and cofactors show promoter-specific differences *in vivo*"

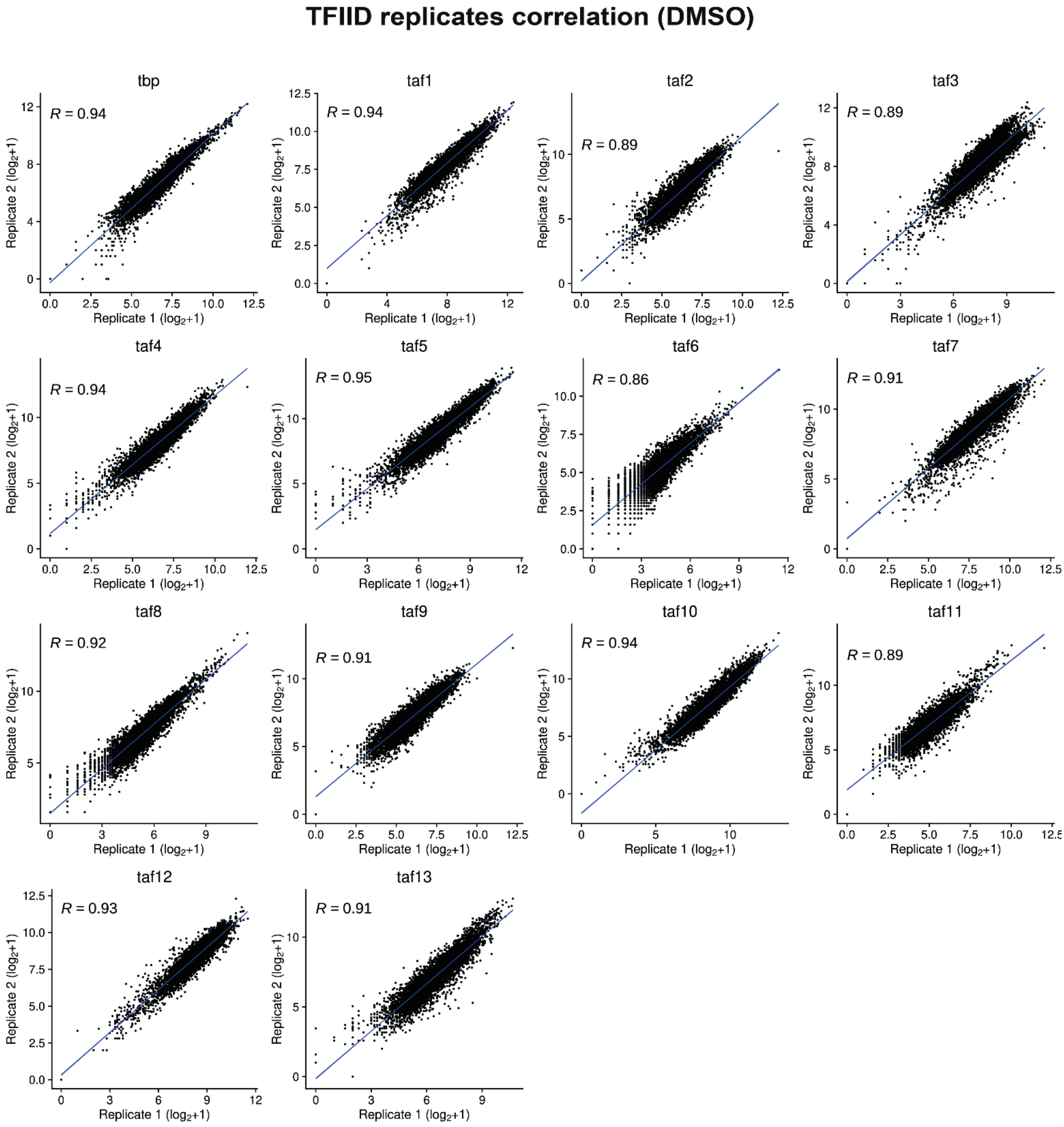


**Figure S1: Pearson correlation of ChIP-nexus read counts between all TFIID subunits replicate experiments performed in DMSO (control) conditions.** Each dot represents the read count calculated within 100 bp around the TSS at an active promoter.

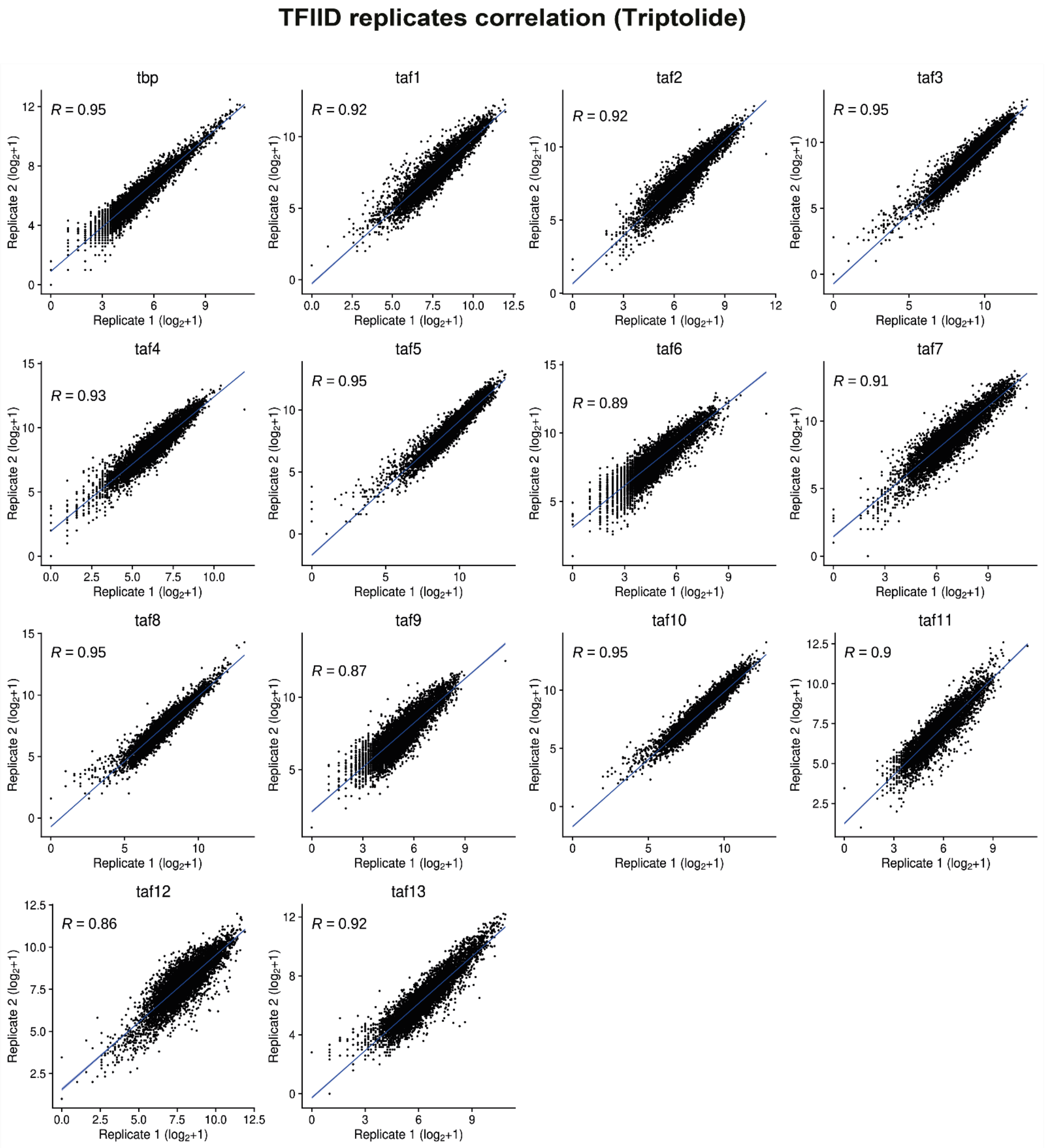


**Figure S2: Pearson correlation of ChIP-nexus read counts between all TFIID subunits replicate experiments performed after triptolide treatment.** Each dot represents the read count calculated within 100 bp around the TSS at an active promoter.

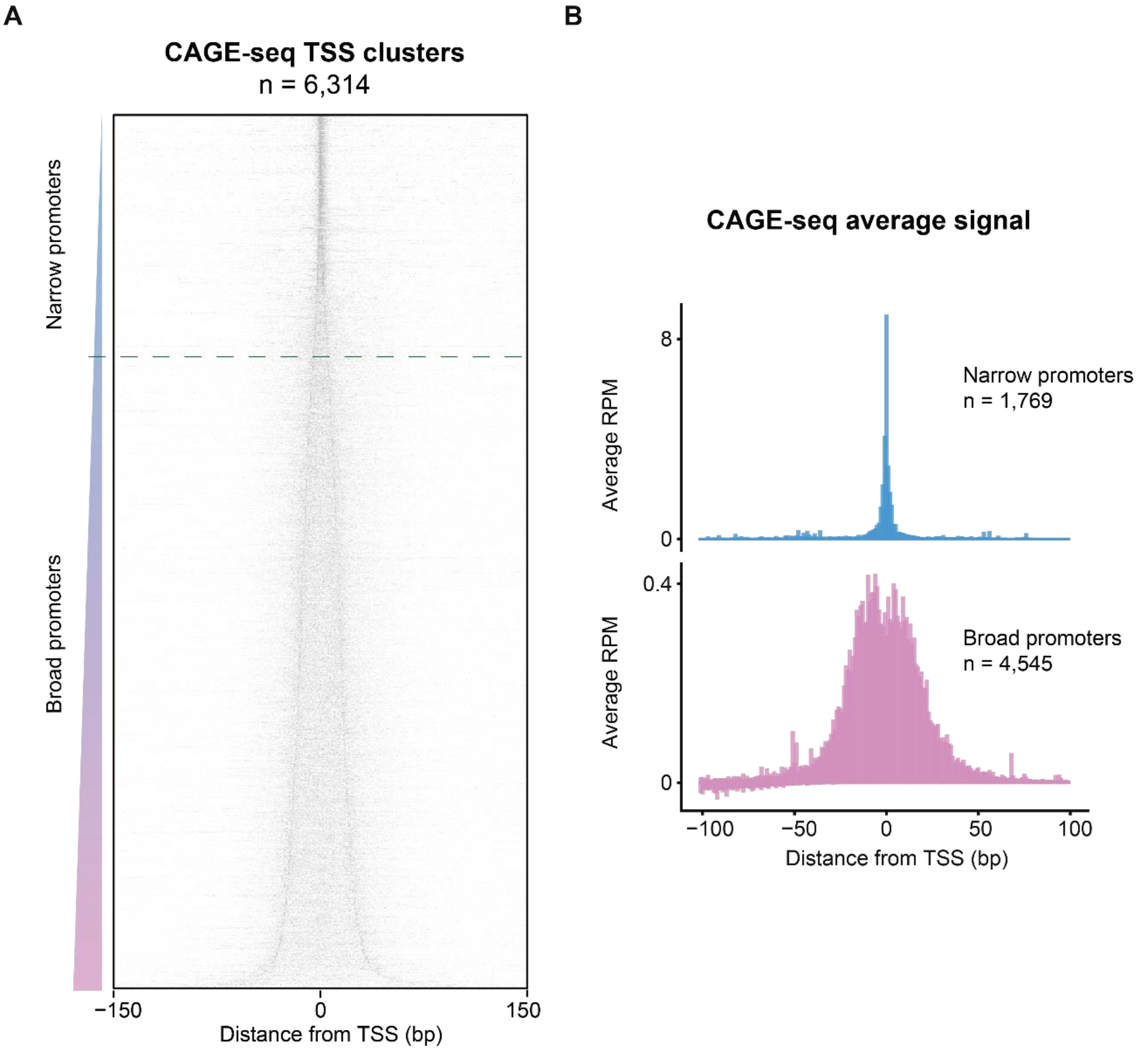


**Figure S3: Selection of narrow TSSs using CAGE-seq data.**
**A)** Heatmap showing the CAGE-seq normalized signal at all the selected active promoters (6314). TSSs are shown centered at the mid position and sorted based on the width of the TSS cluster (from narrower to broader; see Methods). **B)** Average CAGE-seq levels at narrow (cluster width <= 11bp) and broad (cluster width > 11bp) promoters.

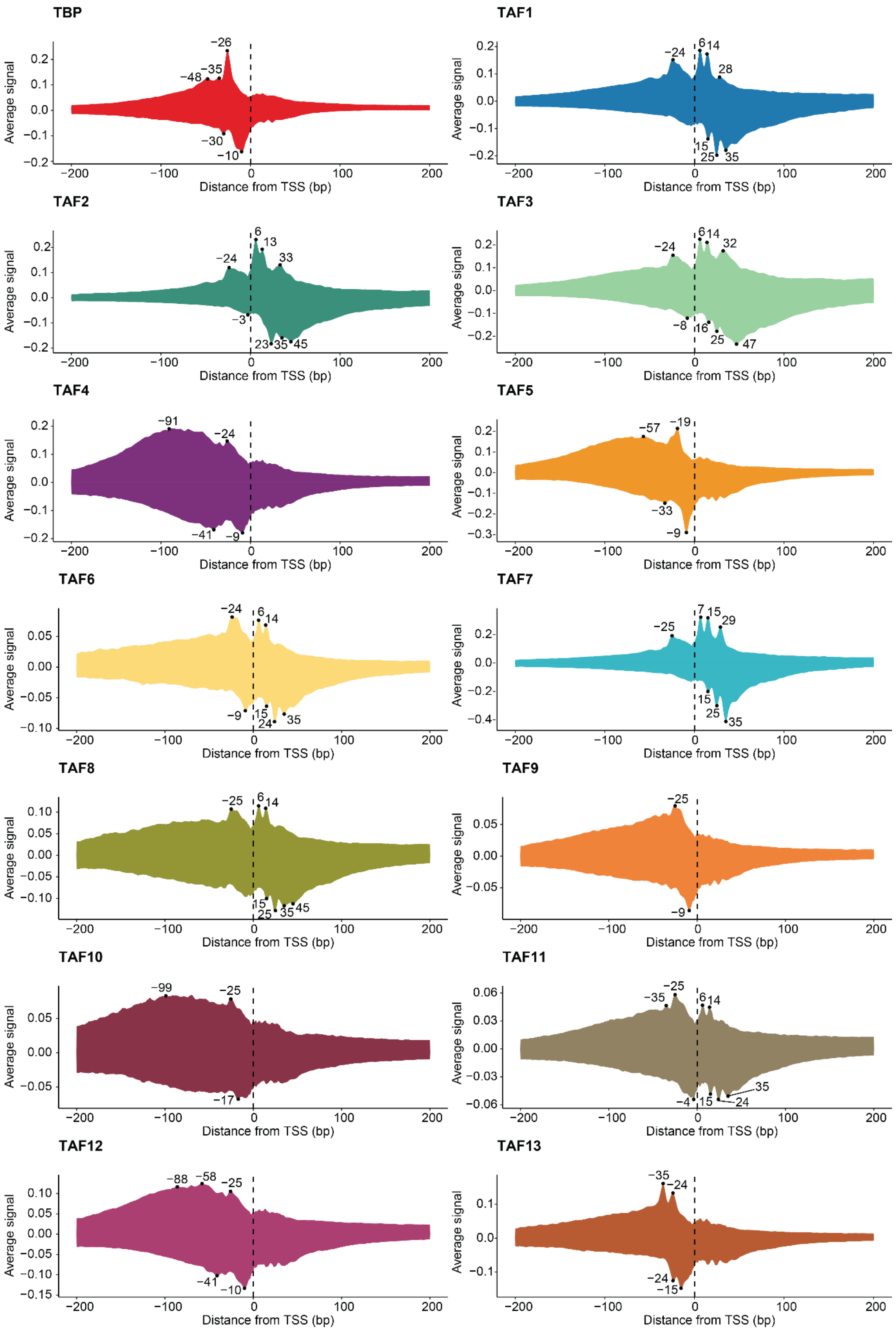


**Figure S4: Average ChIP-nexus binding profile of individual TFIID subunits at narrow promoters**
The average ChIP-nexus binding profile of each TFIID subunit presented in Figure 2 is shown with their corresponding normalized signal. Each profile is centered at the TSS and shown with the binding signal at the positive strand (top line) and negative strand (bottom line) in the same color.

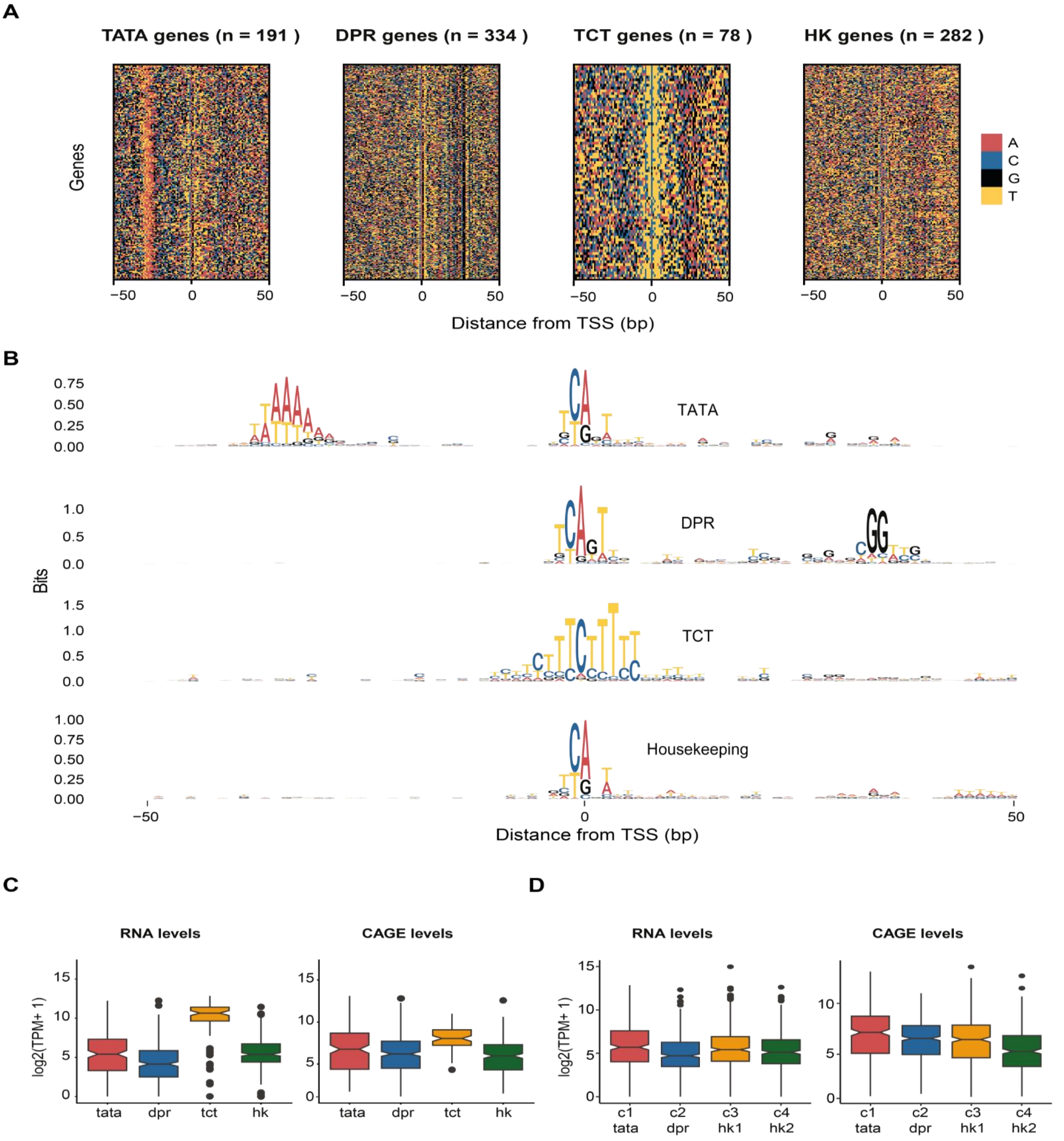


**Figure S5: DNA sequence composition and transcriptional patterns of TATA, DPR TCT and HK promoters.**
**A)** DNA sequence heatmaps showing the differences in DNA composition between TATA, DPR, TCT and HK core promoters. **B)** Position weight matrix (PWM) of the upstream and downstream region of TATA, DPR, TCT and HK core promoters, respectively. **C)** RNA-seq and CAGE-seq mRNA levels at the different promoter classes. **D)** RNA-seq and CAGE-seq mRNA levels at the different clusters in Figure 3.

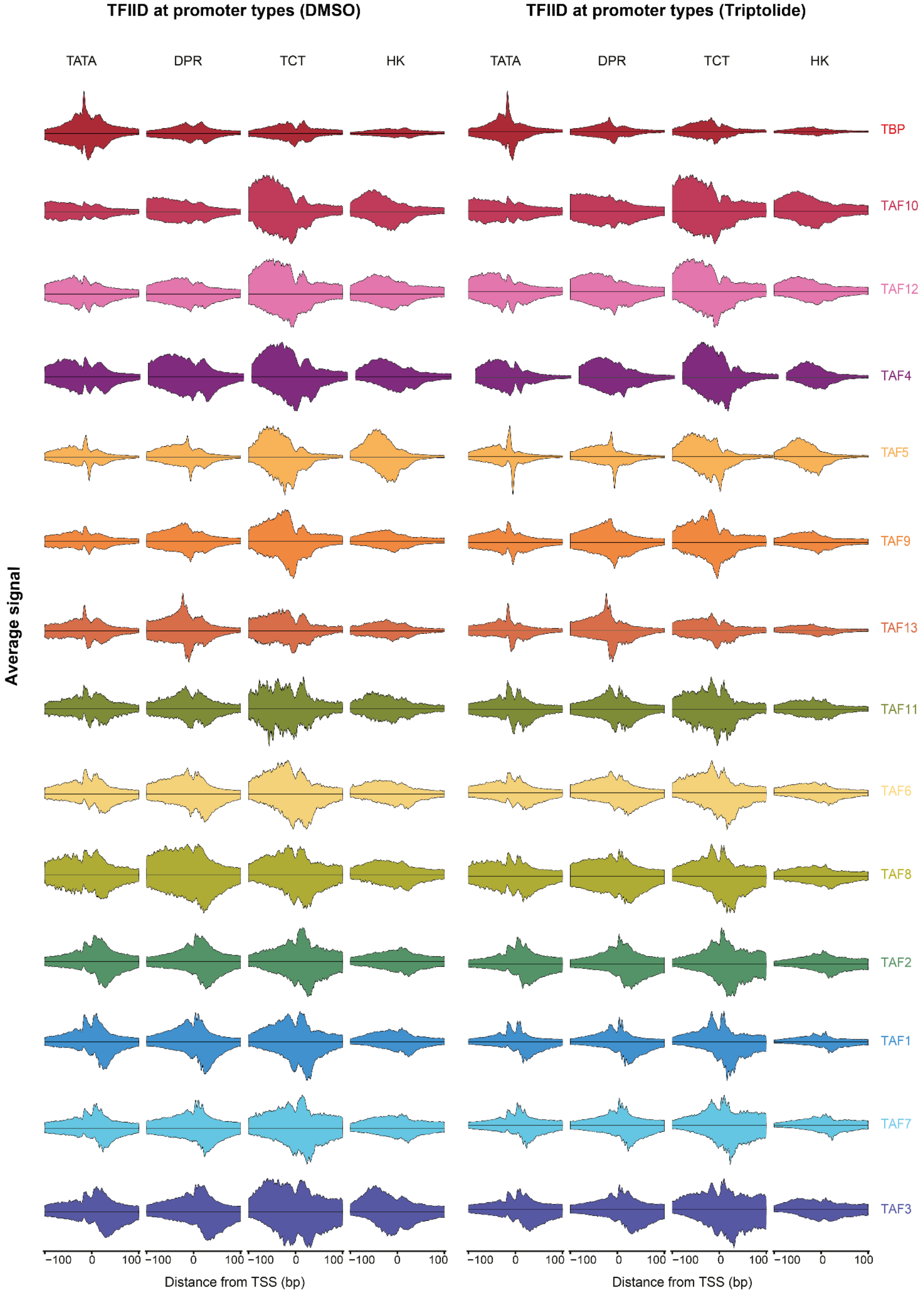


**Figure S6: TFIID binding profile at promoter types.** Average ChIP-nexus binding profile of all TFIID subunits at TATA, DPR, TCT and HK promoters centered at the TSS in DMSO (control) and Triptolide.

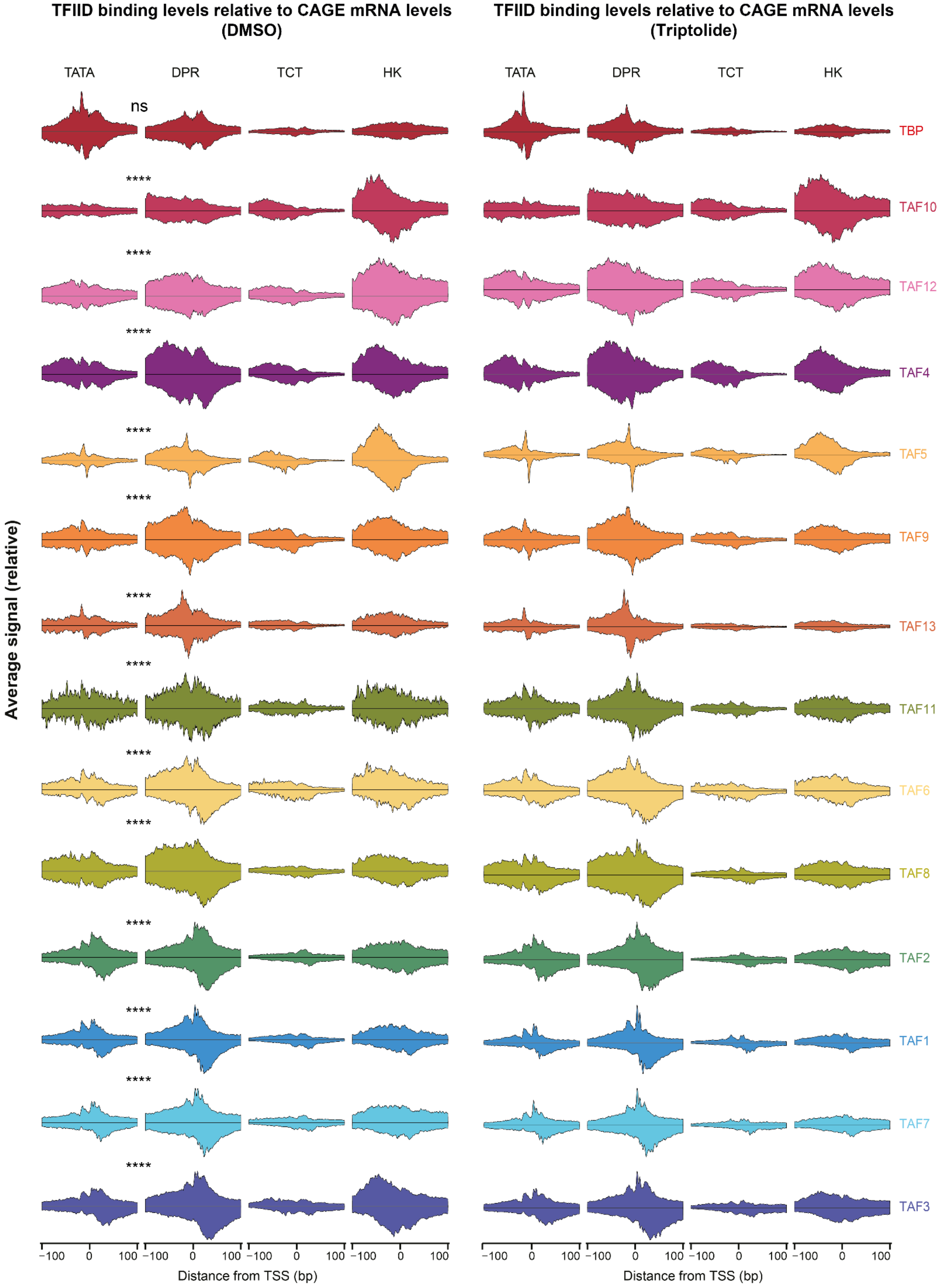


**Figure S7: Expression normalized TFIID binding profile at promoter types.** CAGE-seq normalized average ChIP-nexus binding profile of all TFIID subunits at TATA, DPR, TCT and HK promoters centered at the TSS in DMSO (control) and Triptolide conditions. Significance was calculated comparing the total ChIP-nexus occupancy between TATA and DPR promoters 100bp around the TSS (Wilcoxon test).

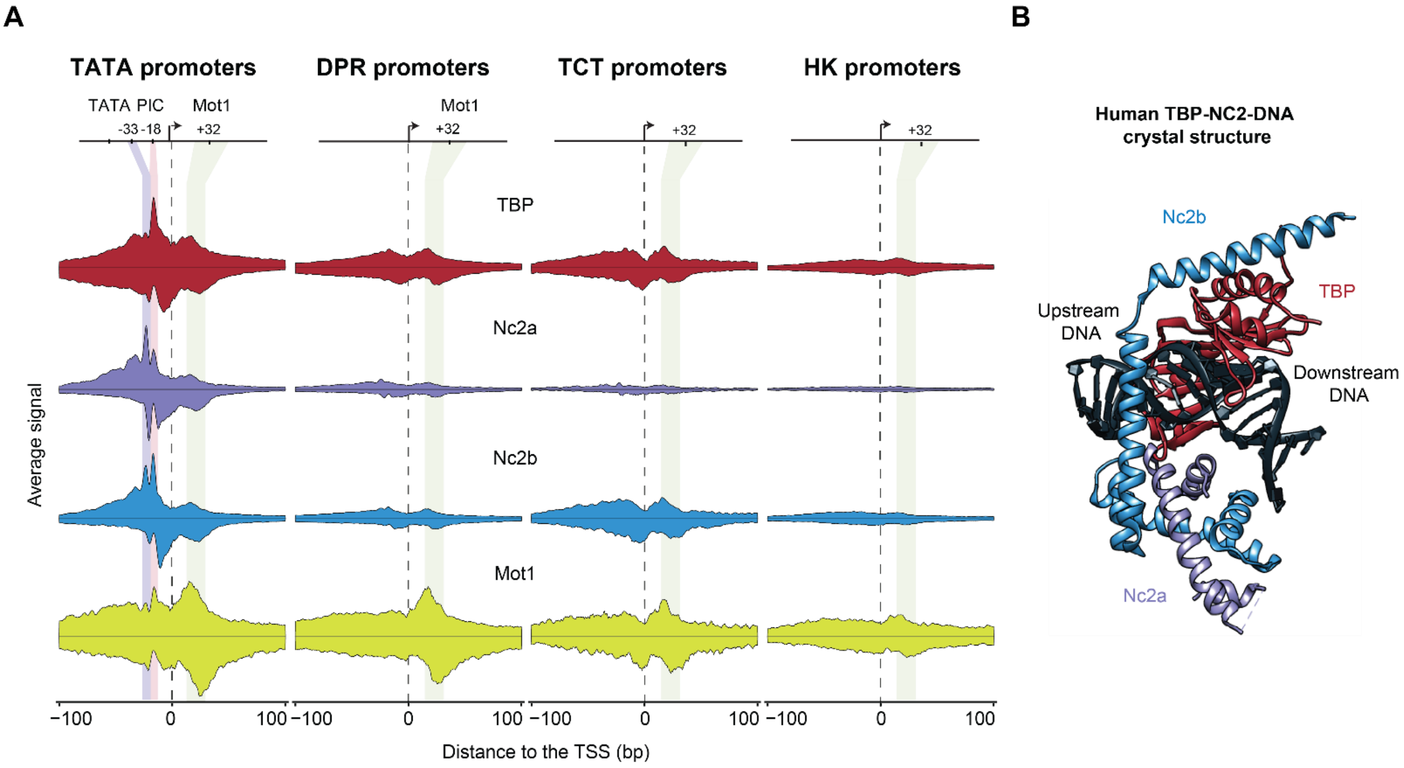


**Figure S8: NC2 and Mo1 binding profile at promoter types.**
**A)** Average Chip-nexus binding profile of all TFIID subunits at TATA, DPR, TCT and HK promoters centered at the TSS. **B)** Crystal structure of human NC2-TBP-DNA.

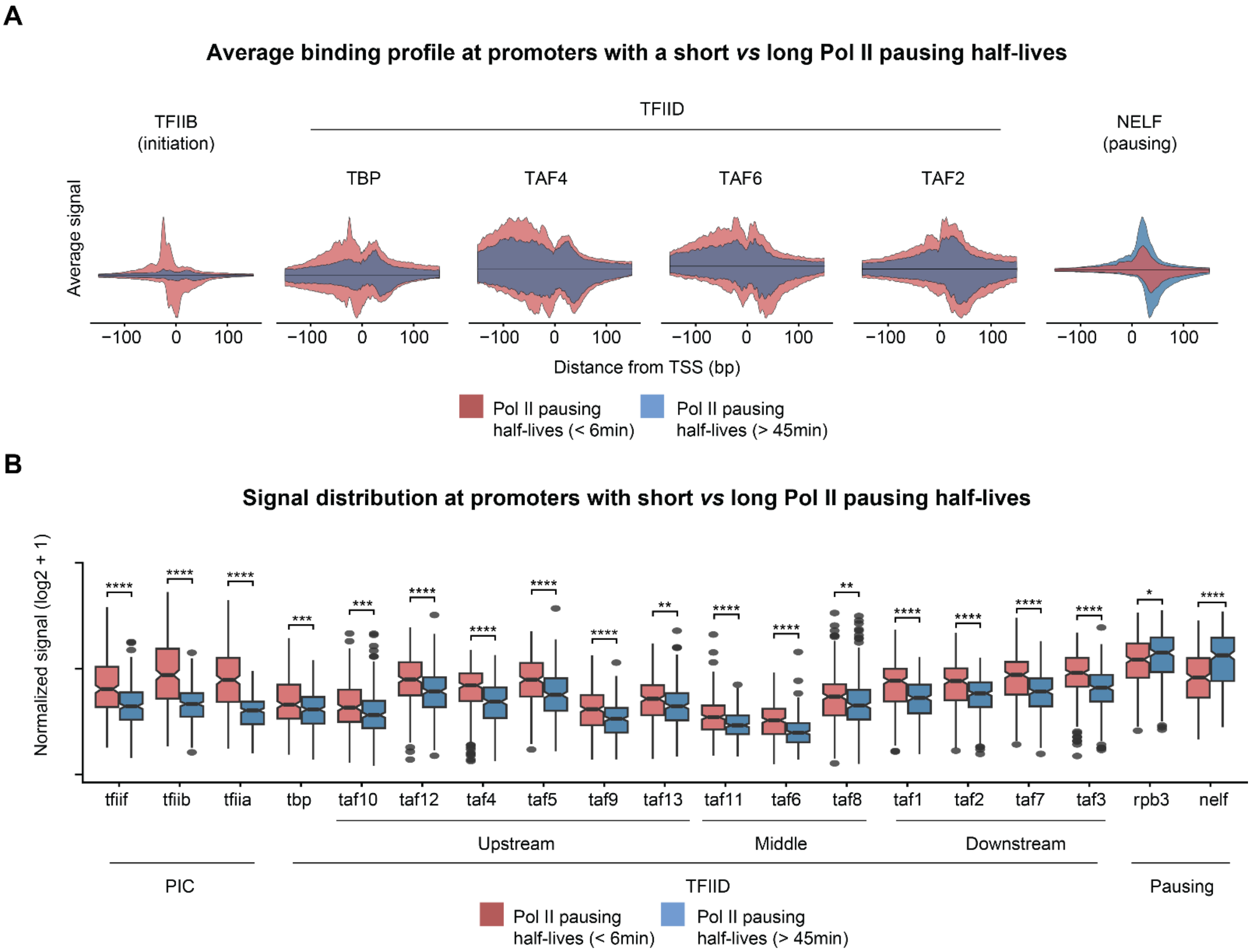


**Figure S9: Downstream TBP and TFIID occupancy is lower at promoters with stably paused Pol II.**
**A)** Average ChIP-nexus binding profile of TFIIB, TBP, TAF4, TAF6, TAF2 and NELF around the TSS of promoters with short (red, half-lives < 6min) and long (blue, half-lives > 45min) Pol II pausing half-lives (see Methods). **B)** Total ChIP-nexus levels at promoters with short (red) and long (blue) Pol II pausing half-lives. Levels of PIC components, including TFIID, are reduced at stably paused promoters, while levels of Pol II and the pausing factor NELF are increased (Wilcoxon test). Binding levels for PIC and TFIID factors are calculated 100bp around the TSS. For TBP, Pol II and NELF ChIP-nexus signal was calculated 50bp downstream of the TSS.
